# MoTRUST: adaptive semantic protection for mosaic single-cell integration and molecular recovery

**DOI:** 10.64898/2026.09.18.752592

**Authors:** Qingchun Liu, Yan Xu

## Abstract

Single-cell multi-omics reveals complementary aspects of cellular identity, but heterogeneous assays produce incomplete modality combinations. Integrating these mosaic measurements while preserving biological structure and recovering missing molecular information remains challenging. We present MoTRUST, combining neural representations, adaptive semantic protection, molecular prediction and conditional residual diffusion. MoTRUST ranked first in overall integration across 30 tasks spanning six scenarios and surpassed the strongest evaluated controls on six of eleven development recovery endpoints. Conditional residual diffusion reduced aggregate marginal prediction scores by 4.4–5.1% across three ATAC-to-RNA data sources while preserving point predictions. Together, these capabilities enable accurate mosaic integration and molecular recovery for multimodal characterization of cellular heterogeneity.

## 1 Introduction

Single-cell multimodal measurements reveal complementary aspects of cellular identity. CITE-seq and REAP-seq jointly profile transcripts and surface proteins using antibody-derived tags (ADT) (Stoeckius et al. 2017; Peterson et al. 2017); sci-CAR and SHARE-seq connect transcription with chromatin accessibility measured by the assay for transposase-accessible chromatin (ATAC) (Cao et al. 2018; Ma et al. 2020). TEA-seq and DOGMA-seq extend profiling to three modalities (Swanson et al. 2021; Mimitou et al. 2021), while scNMT-seq measures additional epigenetic layers (Clark et al. 2018). However, collections assembled across technologies contain different modality combinations, sequencing depths and feature spaces. Within each collection, jointly measured bridge cells connect assays, whereas other cells contribute only their observed modalities. Integrating these mosaic measurements requires reconciling technical heterogeneity while retaining complementary biological information.

Established integration methods offer several strategies. Mutual-nearest-neighbor correction, Harmony and Scanorama align related populations (Haghverdi et al. 2018; Korsunsky et al. 2019; Hie et al. 2019); Seurat provides anchor-based integration and multimodal neighborhoods (Stuart et al. 2019; Hao et al. 2021). LIGER and multi-omics factor analysis separate shared from dataset-specific variation (Welch et al. 2019; Argelaguet et al. 2018, 2020). However, comparisons across distinct molecular feature spaces require additional links. GLUE supplies these through a guidance graph (Cao and Gao 2022), while feature-conversion approaches depend on assumptions about molecular relationships. Balancing correction with biological conservation remains central to integration design and evaluation (Argelaguet et al. 2021; Luecken and Theis 2019; Luecken et al. 2022; Fu et al. 2025; Liu et al. 2025).

Generative models capture modality-specific measurements: scVI models RNA, totalVI combines RNA and protein, and Cobolt connects RNA and accessibility (Lopez et al. 2018; Gayoso et al. 2021; Gong et al. 2021). MultiVI accommodates paired and unpaired trimodal data (Ashuach et al. 2023); scANVI and scArches support annotation and reference mapping (Xu et al. 2021; Lotfollahi et al. 2022). These capabilities still depend on informative cross-modal connections and suitable observation models. Transcriptomic recovery methods such as SAVER, ALRA and MAGIC address noisy measurements (Huang et al. 2018; Linderman et al. 2022; van Dijk et al. 2018), although imputation can alter downstream associations (Hou et al. 2020; Andrews and Hemberg 2019).

Mosaic-specific approaches accommodate heterogeneous modality combinations. scMoMaT uses factorization and feature-derived pseudo-counts, while StabMap maps representations through shared features (Zhang et al. 2023; Ghazanfar et al. 2024). Their linking information remains constrained by proxy measurements or dataset connectivity. MIDAS advances joint integration, imputation and knowledge transfer through information-theoretic disentanglement, and MoNAE couples integration with molecular imputation (He et al. 2024; Tang et al. 2024). Palette addresses imbalanced modality composition through graph-based propagation of latent representations (Sheng et al. 2026). Together, these methods broaden mosaic analysis, but successful alignment or reconstruction alone does not establish local biological preservation or reliable conditional variability. This motivates adaptive control of representation changes alongside explicit molecular prediction and distribution modeling.

We developed MoTRUST, a framework for adaptive integration and molecular recovery from mosaic single-cell multi-omics data. MoTRUST combines gated product-of-experts neural encoding with a spectral alternative and RNA-guided adaptive semantic protection, allowing complementary representations to be integrated while controlling departures from local reference structure. A molecular prediction ensemble recovers missing measurements, while conditional residual diffusion models RNA variation around fixed point predictions (Ho et al. 2020; Song et al. 2021). These complementary outputs connect representation learning with direct evaluation of molecular accuracy and predictive distributions. Across six integration scenarios covering diverse tissues and modality combinations, MoTRUST achieved the highest overall score among the six evaluated methods in all 30 benchmark tasks. Recovery evaluations across paired multi-omics datasets showed advantages on most development endpoints and competitive RNA prediction in held-out donors. Across TEA, BMMC-Multiome and Retina, residual diffusion improved aggregate RNA CRPS over a matched conditional stochastic predictor while retaining the point estimate. Cortical analyses further supported recovery of inhibitory subtype-associated transcriptional signals from accessibility measurements, while paired human–mouse integration demonstrated applicability to species-specific feature spaces. Together, these results establish MoTRUST as an approach for combining mosaic integration, molecular recovery and conditional distribution modeling to characterize cellular heterogeneity across heterogeneous multi-omics collections.

## 2 Results

### 2.1 The MoTRUST model

MoTRUST is a computational framework for integrating mosaic single-cell multi-omics data and recovering unobserved molecular measurements (Fig. 1). Its inputs comprise RNA, chromatin accessibility and protein measurements, together with modality-observation masks and batch identifiers. Jointly measured bridge cells connect modalities, while query cells contribute their available measurements. The framework produces an integrated cell representation, molecular point predictions and conditional RNA samples through complementary integration and recovery branches.

**Figure 1:**
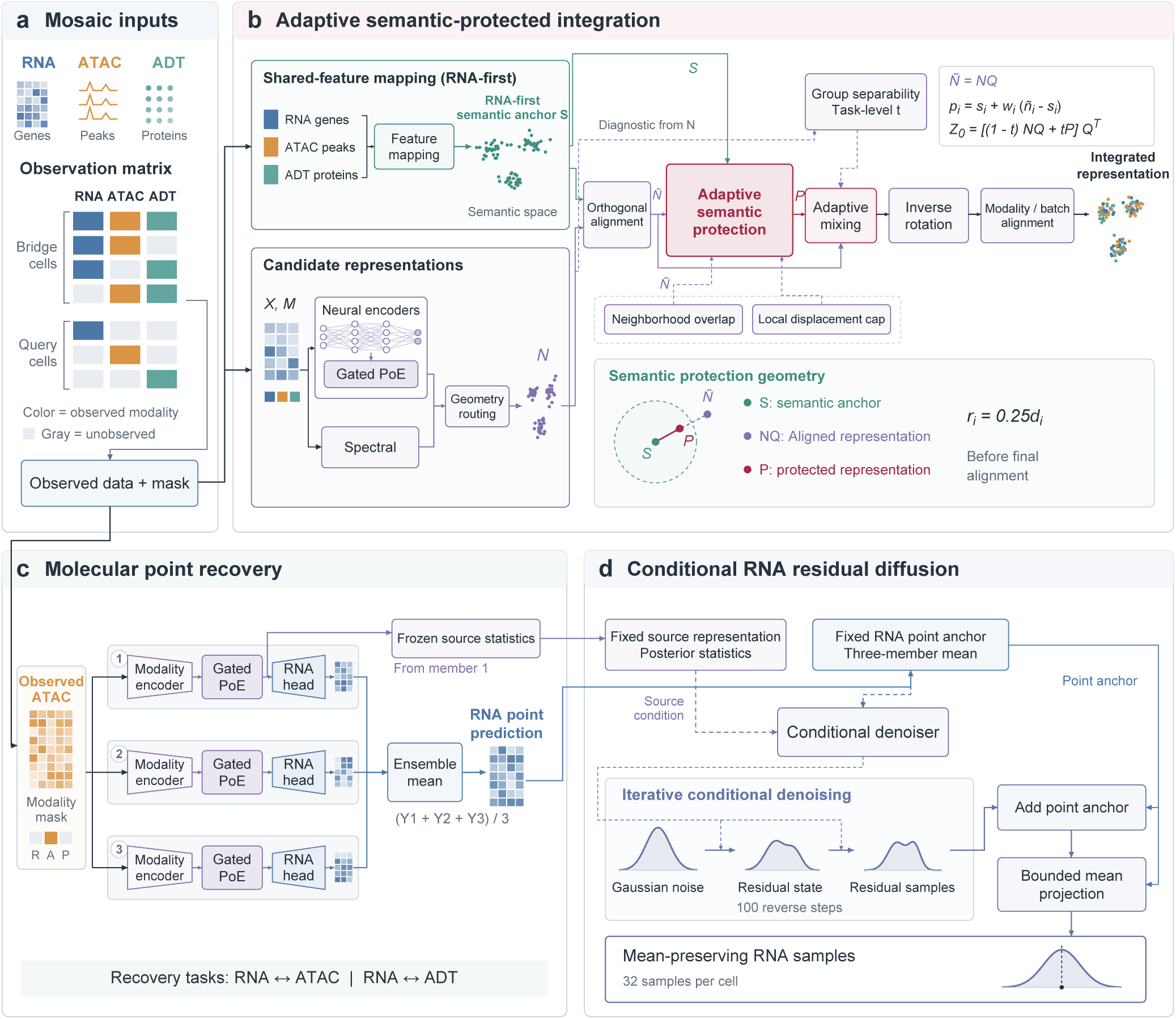
MoTRUST framework for mosaic integration and molecular recovery. a, Observed RNA, ATAC and ADT measurements and their observation mask define bridge and query cells. b, An RNA-first semantic anchor guides protection of an orthogonally aligned neural or spectral candidate. Neighborhood agreement and a local displacement cap determine the retained residual. An unlabeled task-level coefficient mixes protected and aligned representations before inverse rotation and modality/batch alignment. The radius bound applies before final alignment. c, Three independently trained molecular recovery members receive observed source data and masks; their predictions are averaged equally. Recovery and integration are trained separately. In the illustrated ATAC-to-RNA task, Gated PoE combines the observed expert with a Gaussian prior. Posterior statistics from the first fixed member supply the diffusion source condition. d, Conditional RNA residual diffusion uses that source representation and the ensemble point anchor over 100 reverse steps. After adding the point anchor, bounded mean projection yields 32 samples per cell whose empirical mean matches the point prediction to numerical tolerance. Mean preservation does not establish interval calibration. Matrices, embeddings, network icons and distributions are schematic.

For integration, MoTRUST constructs an RNA-guided semantic reference alongside a learned candidate representation. Compatible RNA genes, gene-associated accessibility and mapped proteins are projected into a shared feature space, with observed RNA taking priority when assigning each cell’s semantic anchor. In parallel, neural encoders summarize observed modalities, and a gated product of experts combines their posterior information with a Gaussian prior. Modality gates regulate each expert’s contribution, while the observation mask excludes unavailable measurements. A fixed geometry-based rule selects between this candidate and a spectral alternative before semantic protection (Methods).

Adaptive semantic protection controls how the selected representation is combined with the reference. An orthogonal transformation first aligns their coordinates. Agreement between local neighborhoods determines how much of the candidate’s residual displacement is retained around each semantic anchor, subject to a cap based on the local reference radius. A task-level coefficient derived from observation-group separability then mixes the protected and aligned candidate representations without using cell-type labels. Inverse rotation and subsequent modality and batch alignment yield the integrated representation. The displacement cap applies only to the protected intermediate representation.

Molecular recovery is trained separately from integration to predict target features from observed source modalities. Each of three independently trained members comprises modality encoders, a gated product of experts and target-specific molecular heads. Their predictions are averaged with equal weights, supporting bidirectional RNA–ATAC and RNA–protein recovery. To model variation around RNA point predictions, MoTRUST uses conditional residual diffusion. A denoising network learns to reverse noise added to training residuals, conditioned on fixed posterior statistics from one recovery member and the ensemble RNA prediction. Iterative denoising generates residual samples, which are added to the point prediction and subjected to bounded mean projection. The resulting RNA samples retain the ensemble prediction as their empirical mean while representing conditional variation.

### 2.2 MoTRUST consistently integrates diverse mosaic multi-omics datasets

We first evaluated MoTRUST against five mosaic integration methods: Palette, MIDAS, scMo-MaT, StabMap and MoNAE. The benchmark comprised six scenarios spanning RNA–protein, RNA–accessibility and trimodal measurements: Ab-seq, Retina, two BMMC configurations and two TEA configurations. Five repeats per scenario yielded 30 integration tasks with varying observation patterns and input configurations. All methods were assessed using a common evaluator that combined biological conservation and batch correction into an overall score with weights of 0.6 and 0.4, respectively. Repeats combined task/subsampling variation with neural training seeds; paired comparisons were interpreted descriptively (Methods).

MoTRUST achieved the highest overall score in all 30 tasks (Fig. 2a; Table 1). Its mean score was 0.7258, compared with 0.6753 for Palette and 0.6199 for MIDAS. This advantage extended across all six scenario means, covering both bimodal and trimodal configurations (Fig. 2d). Relative to Palette, the scenario-mean gain ranged from 0.0132 in the second TEA configuration to 0.0766 in the second BMMC configuration, showing that the aggregate ranking was maintained across distinct mosaic settings.

**Figure 2:**
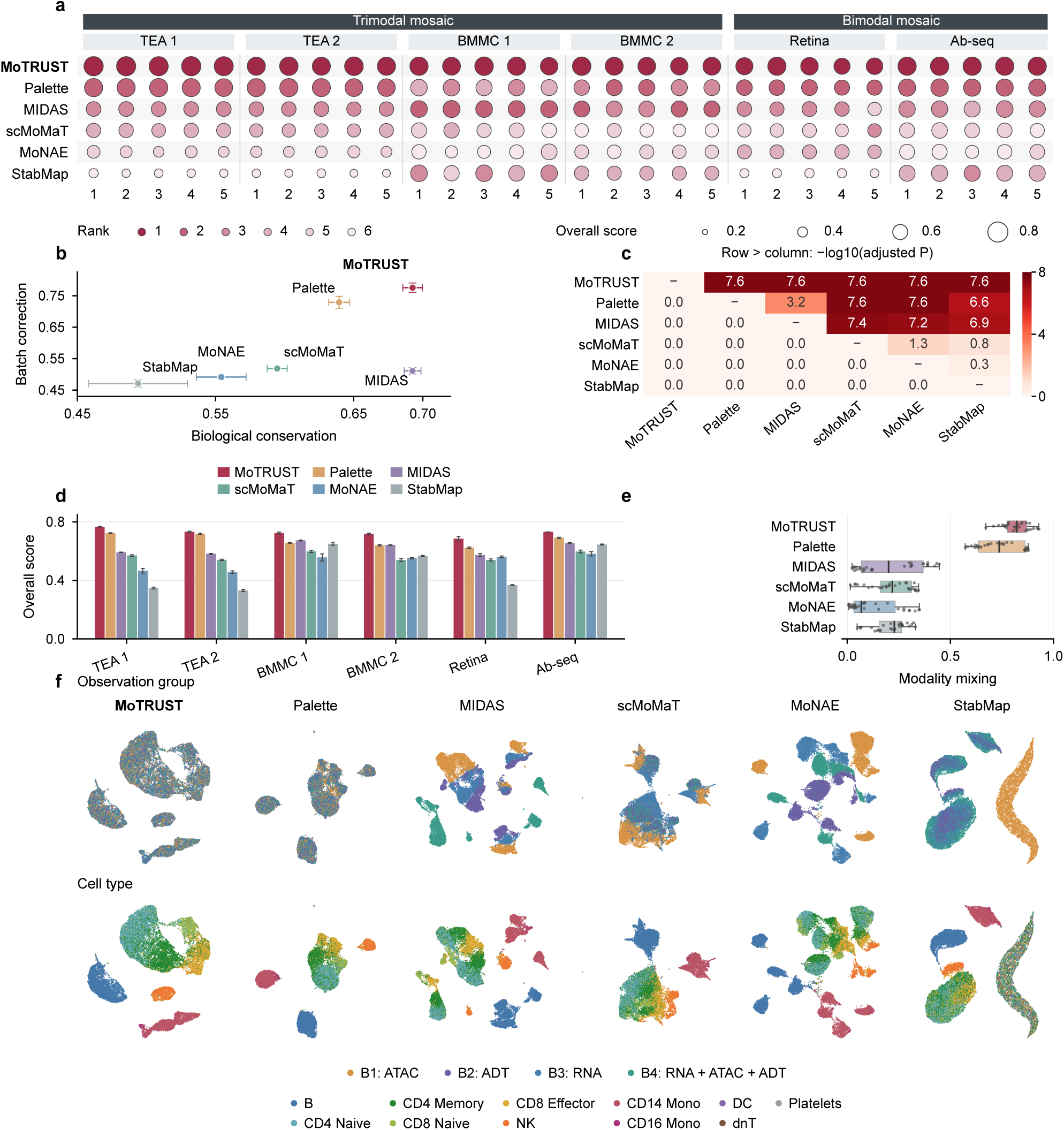
Thirty-task mosaic integration benchmark and integrated representations. a, Overall scores for six methods across 30 tasks. Circle radius is proportional to score; darker color indicates better within-task rank. b, Mean biological conservation and batch correction with SEM across 30 configurations. c, One-sided paired Wilcoxon comparisons with Holm adjustment; numbers and colors show log_10_ adjusted *P* for row-method scores exceeding column-method scores. d, Mean overall scores with SEM across five configurations per scenario. e, Modality-mixing scores: boxes span the interquartile range, center lines mark medians, whiskers extend to values within 1.5 times that range, and dots show all 30 observations. Overall equals 0.6 times biological conservation plus 0.4 times batch correction. f, UMAPs of the same 25,517 cells in the prespecified TEA_s1 random1 task, colored by observation group (top) and 11 annotated cell types (bottom). Native embeddings were standardized and projected independently with identical settings: 15 neighbors, minimum distance 0.3 and seed 2024. Each method uses identical coordinates in both rows, and all panels share one drawing order without subsampling. Task configurations combine sampling and training variability, share source data and are not independent cohorts; paired tests are descriptive.

**Table 1:** Mean integration scores across 30 tasks.

| Method | Overall | Biological conservation | Batch correction |
| --- | --- | --- | --- |
| MoTRUST | 0.7258 | 0.6926 | 0.7755 |
| Palette | 0.6753 | 0.6395 | 0.7289 |
| MIDAS | 0.6199 | 0.6926 | 0.5109 |
| scMoMaT | 0.5643 | 0.5947 | 0.5187 |
| MoNAE | 0.5291 | 0.5542 | 0.4915 |
| StabMap | 0.4848 | 0.4939 | 0.4713 |

The component scores linked this overall advantage to a favorable balance between biological conservation and technical correction (Fig. 2b). MoTRUST improved both components over Palette, with mean conservation scores of 0.6926 versus 0.6395 and correction scores of 0.7755 versus 0.7289. Biological conservation was comparable to MIDAS (0.6926), while batch correction was higher (0.7755 versus 0.5109). Thus, the advantage over MIDAS reflected improved correction at a similar mean conservation level.

MoTRUST also improved mixing across observed modality combinations, with a mean modality-mixing score of 0.8210 compared with 0.7384 for Palette, and ranked first on this measure in 25 of 30 tasks (Fig. 2e). This complementary metric was evaluated separately from the overall score and supported alignment across heterogeneous observation groups. Matched uniform manifold approximation and projection (UMAP) views of the first TEA configuration illustrated observation-group mixing within distinct B-cell, NK-cell and monocyte neighborhoods for MoTRUST (Fig. 2f).

The overall ranking remained stable when the biological-conservation weight varied from 0.40 to 0.80: MoTRUST retained the highest method-mean score at all nine tested weights and led in 28–30 tasks at each weight (Supplemental Fig. S3). Together, these results support consistent overall integration performance across the evaluated mosaic configurations.

### 2.3 Component comparisons reveal task-dependent integration trade-offs

To examine how the integration strategies underlying MoTRUST contributed to its benchmark performance, we compared the full model with four alternatives: unprotected fusion, a semantic-only reference, a fixed 25% candidate residual and local semantic protection without adaptive mixing (Fig. 3a–c).

**Figure 3:**
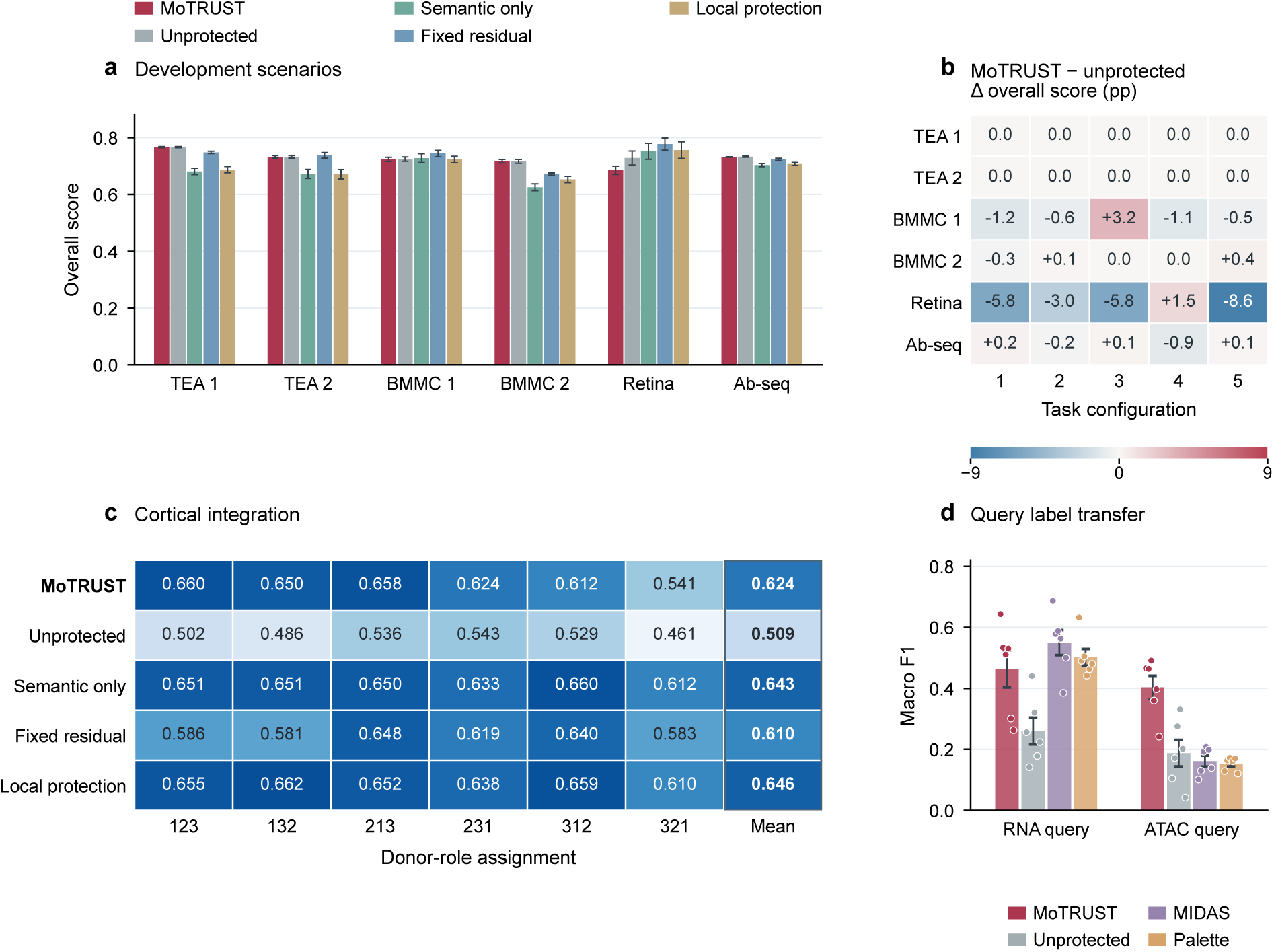
Semantic protection across development and cortical tasks. a, Mean overall scores with SEM across five task configurations per development scenario for each strategy. b, MoTRUST minus unprotected fusion for every matched task, expressed as 100 times the score difference in percentage points (pp). Blue and red indicate negative and positive changes, respectively; numbers identify task configurations within each scenario. c, Overall scores for all six BICCN donor-role assignments and equal-weight means in the outlined column. Each three-digit code gives the donor numbers assigned to bridge, RNA-query and ATAC-query roles, in that order. d, Bridge-based label-transfer macro-F1 for RNA and ATAC queries: bars and error bars show means and SEM across six assignments; dots show every assignment. Task configurations and role assignments share source data and are not independent biological replicates. Strategy configurations differ in intervention and coordinate system, so the comparisons do not isolate each factor.

Across the six benchmark scenarios, MoTRUST exceeded the semantic-only strategy in four scenario means, indicating complementary information beyond the reference in these settings (Fig. 3a). In both TEA scenarios, the task-level rule selected no semantic intervention and preserved the unprotected fusion output. Changes relative to unprotected fusion were below 0.002 in absolute overall score for both BMMC scenarios and Ab-seq, whereas Retina decreased by 0.0431 (Fig. 3b). Thus, introducing protection had little effect in several settings but incurred a larger trade-off in Retina.

The BRAIN Initiative Cell Census Network (BICCN) cortical cohort showed a different pattern. Across six donor-role assignments, MoTRUST improved the mean overall score from 0.5092 with unprotected fusion to 0.6241, with gains in every assignment (Fig. 3c). It also exceeded the fixed-residual strategy (0.6096). However, the semantic-only and local-protection strategies reached higher means of 0.6430 and 0.6458, respectively, showing that retaining more candidate information did not consistently improve on the semantic reference.

These comparisons support a task-dependent balance between reference structure and candidate information. Because the strategies also differed in intervention and coordinate handling, the observed gains characterize complete integration strategies and do not separately establish the contributions of neighborhood agreement, displacement limits or adaptive mixing.

### 2.4 MoTRUST recovers molecular profiles across cohorts and donors

We next evaluated whether MoTRUST could recover unobserved molecular measurements across paired RNA/ATAC datasets (PBMC-10x, Chen-2019 and Ma-2020) and RNA/ADT data (DOGMA). Comparisons included MIDAS and recovery adapters for Palette, StabMap, scMoMaT and MoNAE, using the same target features, evaluation cells and metrics: normalized root mean squared error (RMSE), feature Pearson correlation and ATAC area under the precision–recall curve (AUPRC) (Saito and Rehmsmeier 2015). Palette transferred measured training features, whereas StabMap, scMoMaT and MoNAE representations were coupled to common molecular heads. Each added comparator combined three independently seeded predictions (Methods; Fig. 4).

**Figure 4:**
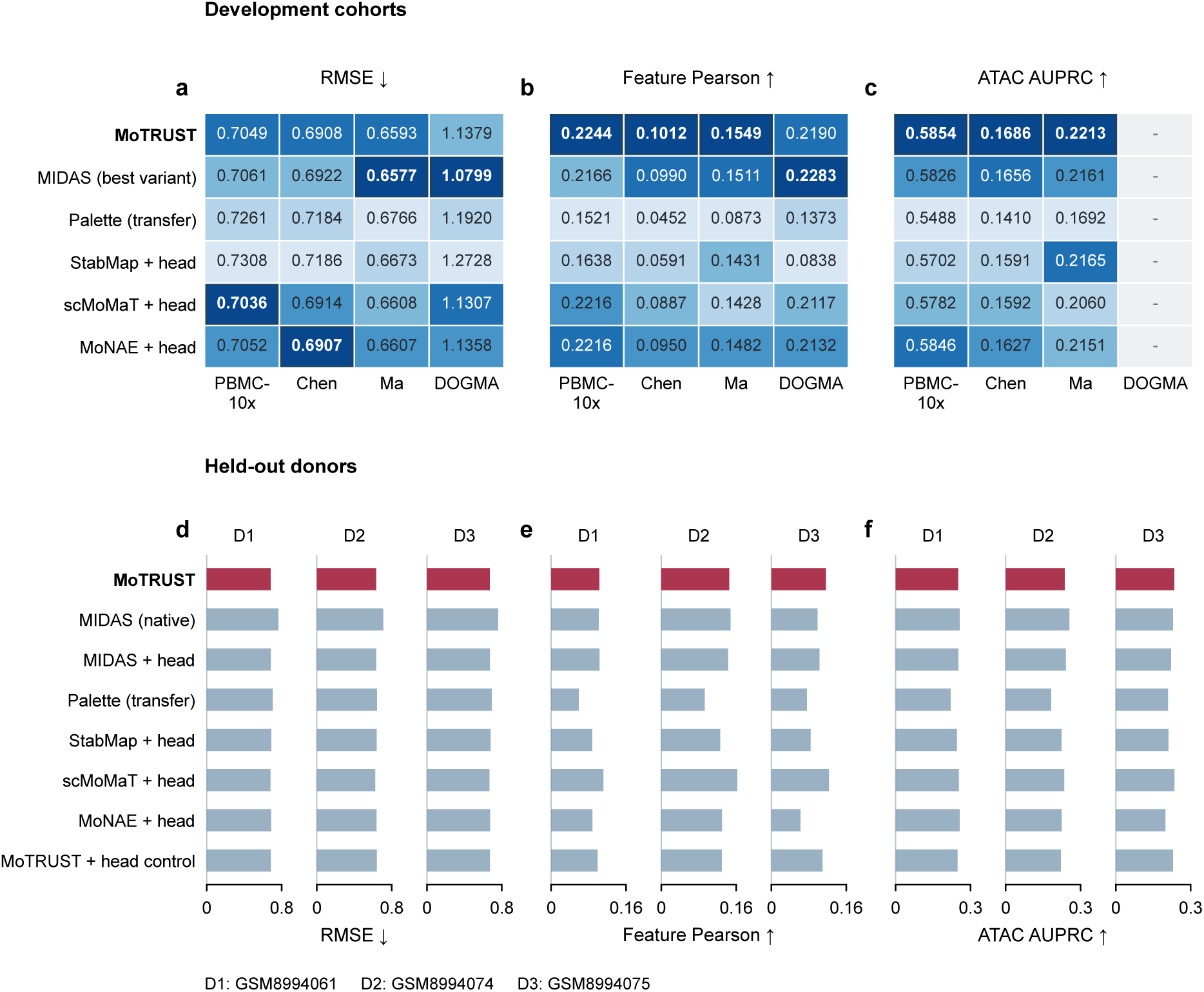
Molecular recovery across six methods. a–c, Scores on development selection partitions. Numbers show actual scores; darker cells indicate better within-cohort ranks and outlined bold values mark the best. Arrows indicate the direction of better performance. Chen and Ma denote Chen-2019 and Ma-2020. MIDAS uses the strongest evaluated variant per endpoint. DOGMA has no ATAC target. d–f, Scores for three held-out GSE297529 donors, shown in separate tracks with identical zero-based scales within each panel. Each bar represents one donor’s aggregate score under the specified protocol. RMSE and feature Pearson average both directions; AUPRC uses ATAC targets only. “+ head” denotes a common molecular-head adapter; Palette uses feature transfer, including the declared PCA fallback for all three Chen-2019 ATAC-to-RNA fits. Added adapters allow source-only query data during representation fitting, unlike fixed-model MoTRUST/MIDAS inference. Endpoint-specific MIDAS variants are listed in Supplemental Table S9. These point-prediction comparisons do not measure diffusion’s contribution.

On the development selection partitions, MoTRUST achieved the highest feature correlation and ATAC AUPRC in all three RNA/ATAC datasets, leading on six of eleven cohort–metric endpoints (Fig. 4a–c). In PBMC-10x, its correlation was 0.2244 versus 0.2216 for the strongest comparator, MoNAE with a molecular head. In Chen-2019, ATAC AUPRC was 0.1686 versus 0.1656 for the strongest MIDAS control. RMSE differences were small in PBMC-10x and Chen-2019, where scMoMaT and MoNAE heads, respectively, performed best. MIDAS variants achieved the lowest RMSE in Ma-2020 and DOGMA and the highest correlation in DOGMA. Thus, the clearest developmental advantage was recovery of feature-level associations and accessible regions in the evaluated RNA/ATAC settings.

Across three held-out GSE297529 donors, MoTRUST outperformed Palette, StabMap and MoNAE adapters on all three equal-donor summary metrics (Fig. 4d–f). However, scMoMaT with a molecular head achieved lower RMSE (0.6571 versus 0.6629), higher correlation (0.1322 versus 0.1216) and slightly higher ATAC AUPRC (0.2410 versus 0.2405). Native MIDAS retained the highest ATAC AUPRC (0.2469). The added adapters used source-only query observations during representation fitting, whereas MoTRUST and MIDAS used fixed-model inference; these results therefore compare the stated deployment settings rather than a uniformly inductive protocol.

Together, these comparisons support competitive molecular recovery with consistent correlation and accessibility advantages in the development RNA/ATAC cohorts, while showing that rankings depend on cohort and metric. Conditional residual diffusion was evaluated separately from these point predictions.

### 2.5 Conditional diffusion improves aggregate RNA prediction scores across three data sources

We next asked whether conditional residual diffusion could improve RNA predictive distributions while retaining the molecular point estimate. We evaluated ATAC-to-RNA prediction in TEA, BMMC-Multiome and Retina, extending recovery evaluation to three data sources used in the integration benchmark. Within each source, diffusion and a conditional stochastic head shared a fixed point anchor, source coordinates, training budget and bounded mean projection. Evaluation used 18,619 held-out cells across the three sources, with 2,000 RNA features and 32 predictive samples per cell (Methods).

Diffusion reduced aggregate continuous ranked probability score (CRPS), averaged over all evaluated cells and RNA features, in all three sources. Relative to the conditional head, mean CRPS decreased by 4.86% in TEA, 4.41% in BMMC-Multiome and 5.07% in Retina (Supplemental Fig. S1; Supplemental Table S1). CRPS evaluates agreement between a marginal predictive distribution and its observed target, with lower values indicating better predictions (Gneiting and Raftery 2007). All three matched training seeds and all nine cross-seed pairings per source favored diffusion (Supplemental Table S8). These repeated fits support a consistent ordering on each fixed split. The sampled mean retained the point prediction to within 1.91 × 10*^−^*^6^, as enforced by projection.

Thus, conditional residual diffusion added a reproducible aggregate marginal-scoring benefit across these three ATAC-to-RNA settings without changing the point estimate. This evidence concerns within-library cell holdouts; improvements were not uniform across expression strata or joint-distribution diagnostics (Supplemental Fig. S4).

### 2.6 MoTRUST supports cortical integration and cell-type transfer

To test integration in a cortical cohort, we analyzed 9,000 BICCN cells from three donors under six assignments of bridge, RNA-query and ATAC-query roles (BRAIN Initiative Cell Census Network (BICCN) 2021). Only observed query modalities entered integration (Fig. 5a–c). MoTRUST achieved a mean overall score of 0.6241, compared with 0.5492 for Palette and 0.5473 for MIDAS, and exceeded both comparators in every assignment (Fig. 5d). Relative to MIDAS, this advantage combined similar biological conservation (0.5706 versus 0.5708) with higher batch correction (0.7043 versus 0.5120).

**Figure 5:**
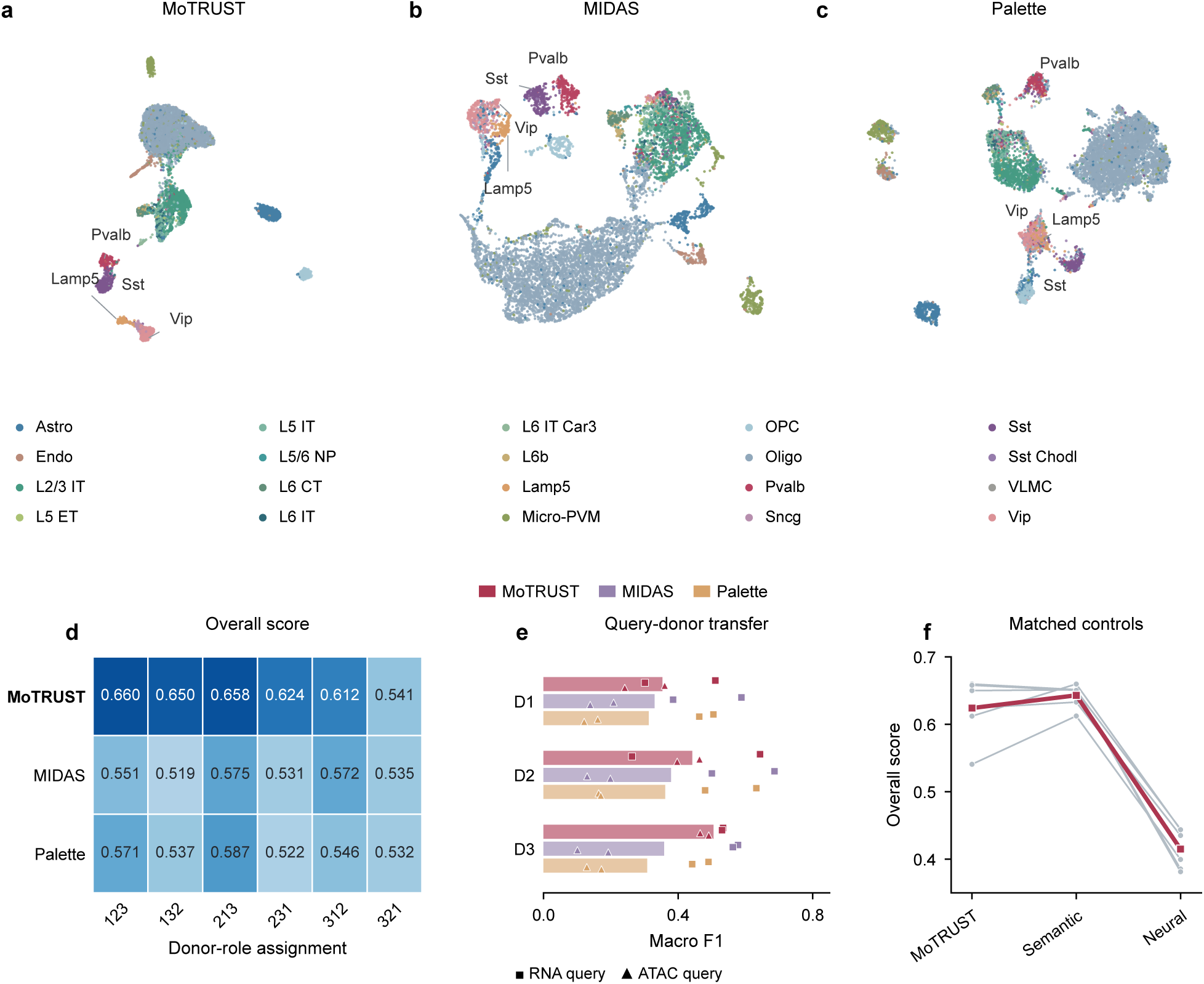
Cortical external-cohort evaluation. a–c, Fixed UMAP coordinates (McInnes et al. 2018) for all 9,000 cells in the first donor-role task (15 neighbors, minimum distance 0.3, seed 2024). Colors identify all 20 cell types; direct labels locate Lamp5, Pvalb, Sst and Vip. d, Numeric overall scores for each of six role assignments. e, Label transfer from 15 bridge neighbors. Bars show mean macro-F1 by donor and method; squares and triangles show all RNA-query and ATAC-query observations, respectively, with four observations per donor-method pair. f, Matched semantic and neural controls. Gray lines connect the scores of the same role assignment across configurations; the red line connects equal-weight means. All six tasks share the same cells from three donors and are not independent cohorts. Only observed query modalities participate in unsupervised integration.

Label transfer showed a pronounced advantage for ATAC queries. MoTRUST reached a mean macro-F1 of 0.4032, compared with 0.1614 for MIDAS and 0.1528 for Palette; for RNA queries, its macro-F1 was 0.4638, below MIDAS (0.5501) and Palette (0.5020). Averaging query roles within each donor, MoTRUST exceeded both comparators in all three donors (Fig. 5e). The matched semantic-only control retained a higher overall integration score (Fig. 5f), consistent with the component comparison in Section 2.3. Together, these results extend the integration advantage over MIDAS and Palette to the cortical cohort, with the clearest label-transfer gain in cells lacking observed RNA.

### 2.7 MoTRUST recovers inhibitory subtype markers from accessibility

We next examined whether cortical integration supported recovery of molecular signals associated with inhibitory subtypes. Measured RNA and gene-local accessibility provided a reference across cortical subclasses (Fig. 6a–e). We focused on PVALB, SST, VIP and LAMP5, established markers of inhibitory diversity (Scala et al. 2021; DeFelipe et al. 2013). Within inhibitory cells, their donor-mean RNA marker AUROCs ranged from 0.8808 to 0.9329, whereas accessibility specificity depended on the marker and genomic window (Fig. 6c).

**Figure 6:**
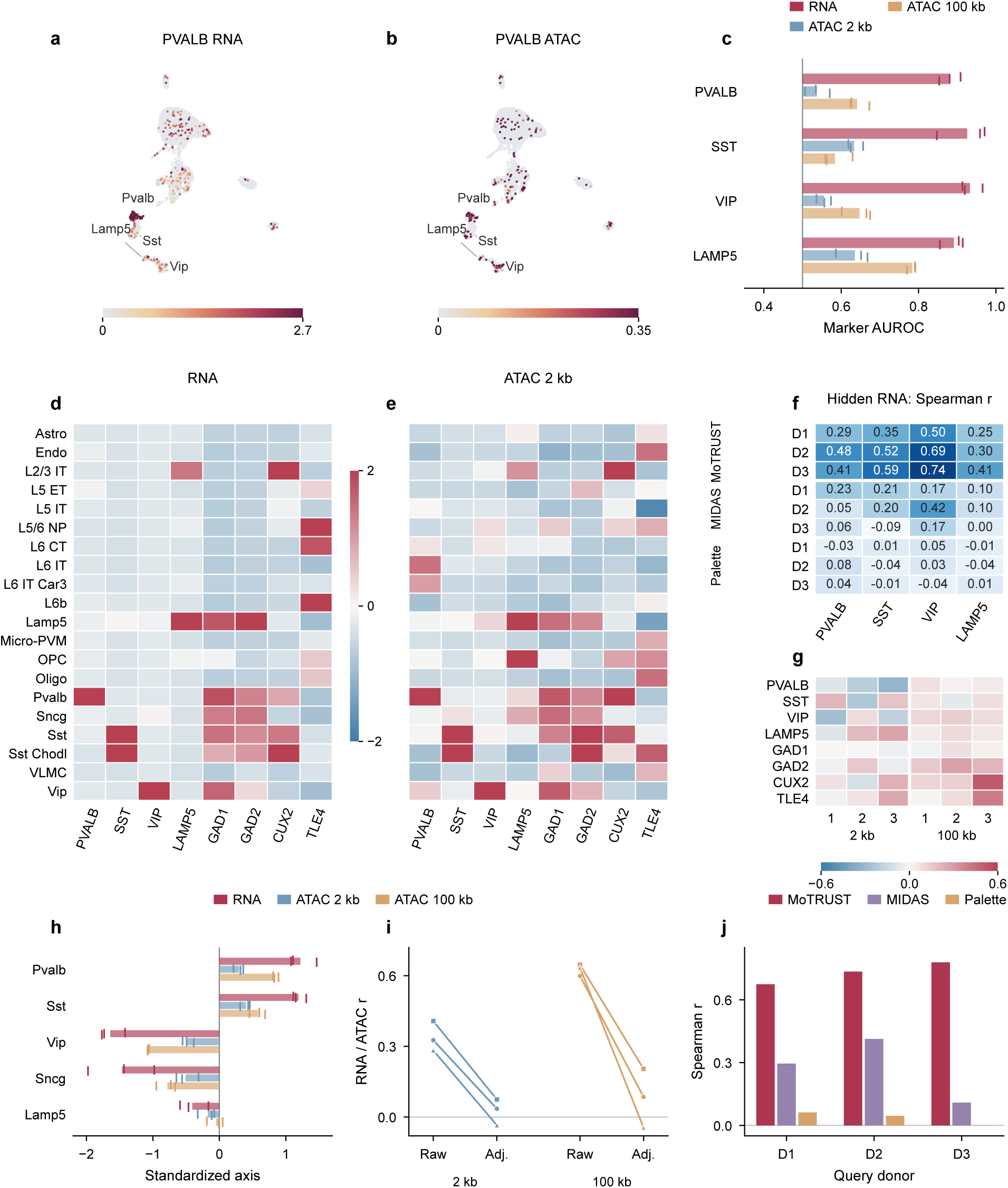
Inhibitory subtype markers and measured cross-modal evidence. a,b, Measured PVALB RNA and 2-kb ATAC (log1p-normalized counts) over the same fixed UMAP; each color scale ends at the signal’s 99th percentile. c, Marker AUROC for RNA and two ATAC windows. Bars extend from chance (0.5) to donor means; ticks show all three donors. d,e, Donor-equal RNA and 2-kb ATAC subtype profiles, standardized within each gene. Colors saturate beyond *z* = 2. f, Hidden-RNA marker recovery in inhibitory cells: numbers and colors show each donor’s Spearman correlation after averaging role assignments. g, RNA/ATAC residual Spearman correlations after subtype and depth adjustment, for both windows and all three donors. h–j, Secondary four-gene axis: signed subtype means with three donor ticks (h), donor-paired measured associations before and after adjustment (i), and hidden-RNA recovery for each donor (j). In i, circles, squares and triangles identify D1, D2 and D3, respectively. Each bar in j is one donor mean over role assignments. The axis is 0.5[*z*(LHX6) + *z*(SOX6) *z*(PROX1) *z*(NR2F2)]. Marker expression and local accessibility do not establish transcription-factor binding or causal regulation.

With query RNA hidden, the mean RNA signal of 15 bridge neighbors recovered each marker from the integrated representation. This readout evaluates the embedding separately from the molecular prediction heads and residual diffusion. Donor-equally weighted Spearman correlations were 0.3954 for PVALB, 0.4852 for SST, 0.6435 for VIP and 0.3213 for LAMP5, exceeding MIDAS and Palette for every marker (Fig. 6f). Restricting evaluation to inhibitory cells reduced the contribution of broad excitatory–inhibitory composition to these associations.

A complementary four-gene axis combining LHX6, SOX6, PROX1 and NR2F2 distinguished Pvalb/Sst-associated from Vip/Sncg-associated profiles in measured RNA and accessibility (Fig. 6h). Its hidden-RNA recovery correlation was 0.7294, compared with 0.2723 for MIDAS and 0.0358 for Palette, while the matched semantic-only control reached 0.7466 (Fig. 6j). Adjusted gene-level associations varied across markers and windows (Fig. 6g); RNA–ATAC agreement for the four-gene axis also attenuated after accounting for subtype and molecular depth (Fig. 6i). These results support recovery of subtype-associated transcriptional signals, without establishing regulatory activity within subtypes.

### 2.8 MoTRUST integrates human and mouse cells across feature spaces

We next extended the analysis to human and mouse cells, where conserved cell identities coexist with species-specific molecular features (Hodge et al. 2019). The paired MOp subset comprised 3,000 cells per species, with 4,659 shared homologous RNA groups and 5,000 ATAC peaks per species. A common RNA semantic anchor connected the species, while separate ATAC decoding heads accommodated their distinct accessibility features.

MoTRUST achieved the highest overall score among the six evaluated methods: 0.7503, compared with 0.7093 for Palette and 0.6682 for MIDAS (Table 2). Its species/batch mixing score was also highest. The overall ranking remained first across all nine tested biological-conservation weights from 0.40 to 0.80 (Supplemental Fig. S3).

**Table 2:** Human–mouse MOp comparison at one fixed training seed.

| Method | Overall | Biology | Mixing | Macro-F1 |
| --- | --- | --- | --- | --- |
| MoTRUST | 0.7503 | 0.6480 | 0.9038 | 0.5869 |
| Palette | 0.7093 | 0.6717 | 0.7658 | 0.5495 |
| MIDAS | 0.6682 | 0.7055 | 0.6121 | 0.6906 |
| scMoMaT | 0.4815 | 0.4009 | 0.6024 | 0.1078 |
| StabMap | 0.3427 | 0.3721 | 0.2986 | 0.0800 |
| MoNAE | 0.5222 | 0.5587 | 0.4674 | 0.2455 |
| RNA anchor | 0.6052 | 0.7051 | 0.4552 | 0.4740 |

Bidirectional label-transfer macro-F1 reached 0.5869, exceeding Palette (0.5495) but remaining below MIDAS (0.6906), which also led biological conservation. Relative to the RNA-anchor control, MoTRUST likewise combined higher mixing with lower conservation (Table 2). These results support integration across the evaluated species-specific feature spaces, with the overall advantage driven by stronger mixing. The evidence is limited to one paired subset and one training seed, with species confounded with assay batch; ATAC-only cross-species transfer was not tested.

## 3 Discussion

MoTRUST provides a framework for combining mosaic integration with molecular recovery while controlling how complementary measurements reshape a cell representation. Across heterogeneous modality configurations, adaptive semantic protection supported a favorable balance between biological conservation and technical correction. Molecular prediction ensembles and conditional residual diffusion extended this representation-level capability to missing measurements and RNA predictive distributions. Together, these results show how integration, point recovery and distribution modeling can be connected within one framework and assessed against their respective biological and statistical objectives.

This design builds on complementary approaches to mosaic analysis. MIDAS uses generative modeling and information-theoretic disentanglement to integrate measurements and recover missing modalities (He et al. 2024), whereas Palette propagates representations through modality connections to accommodate imbalanced observation patterns (Sheng et al. 2026). MoTRUST adds an explicit, locally constrained intermediate representation: neighborhood agreement regulates candidate information around an RNA-guided anchor, and observation-group separability determines the strength of intervention. The component comparisons indicate that the value of this intervention depends on the input configuration. They support adaptive use of complementary representations, while leaving the effects of individual protection components unresolved.

The cortical and cross-species analyses illustrate how this balance relates to downstream use. Cortical integration supported transfer to ATAC-only queries and recovery of inhibitory subtype-associated RNA signals, connecting alignment quality to interpretable molecular read-outs. In human–mouse integration, shared RNA coordinates connected species-specific accessibility spaces. However, stronger mixing did not uniformly improve cell-type transfer or conservation: MIDAS retained advantages in several such comparisons, and the semantic-only control remained strong in the cortical cohort. Thus, an integrated representation is most informative when evaluated for the intended biological question as well as for aggregate alignment quality.

The recovery branch provides a complementary distinction between estimating molecular abundance and representing conditional variation. Averaging independently trained predictors follows the ensemble principle of combining complementary prediction errors (Lakshminarayanan et al. 2017). Conditional residual diffusion then models variation around the fixed ensemble estimate. Its aggregate CRPS improvement across TEA, BMMC-Multiome and Retina therefore reflects added marginal-distribution value without a change in point accuracy. The gains were concentrated among zero-valued observations, with higher nonzero-target CRPS and no consistent energy-score improvement (Supplemental Fig. S4). This supports aggregate marginal prediction within the evaluated settings, while covariance recovery and reliable intervals remain separate objectives. The donor-level coverage diagnostics similarly show that empirical calibration requires direct assessment beyond point error or ensemble agreement (Guo et al. 2017; Lei et al. 2018).

The present scope is shaped by reference coverage and evaluation design. RNA-first anchoring and gene-proximity mapping provide useful cross-modal links, but can limit representations of poorly mapped features or states absent from bridge cells; low rare-type recall identifies one practical consequence. The integration tasks reuse source datasets, the diffusion tests hold out cells within libraries, and the cross-species experiment uses one paired subset and one training seed. In addition, recovery rankings depend on the cohort, readout and use of source-only query observations by comparator adapters. These conditions define the current validation envelope. Extending distributional evaluation to unseen donors and improving calibration under molecular sparsity would clarify when conditional samples support biological analyses beyond the aggregate prediction score.

MoTRUST brings adaptive representation control, molecular point recovery and conditional RNA sampling into a common analytical framework. Its results support mosaic analysis across the evaluated tissues, modality combinations and species-specific feature spaces. By making each output explicit, the framework enables users to assess the contribution most relevant to their task, from constructing integrated cell maps to recovering missing molecular profiles.

## 4 Methods

### 4.1 Overview

MoTRUST takes molecular measurements, batch identities and an observed-modality mask and produces integrated cell representations, molecular point predictions and conditional RNA samples (Fig. 1). Integration combines an RNA-guided semantic anchor with a neural or spectral candidate through adaptive semantic protection and subsequent modality/batch alignment. A separately trained recovery model predicts unobserved RNA, ATAC or ADT from available source modalities. Conditional residual diffusion extends ATAC-to-RNA recovery by sampling around a fixed RNA point estimate. Biological annotations are reserved for evaluation; unavailable measurements contribute neither encoder information nor reconstruction targets.

### 4.2 Observed-modality encoding and candidate representations

For cell *i* and modality *m*, let *x_im_* denote the measurements and *M_im_* ∈ {0, 1} indicate observation. Each encoder yields diagonal Gaussian parameters *µ_im_* and *ℓ_im_* (log variance). A modality-specific network maps their concatenation, detached from the encoder gradient, to a gate *g_im_* = 2 sigmoid(·); it contains a 64-unit ReLU layer and an output layer initialized to give unit gates. Including a unit Gaussian prior, gated product-of-experts fusion gives elementwise precision and mean

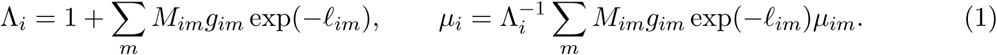

The posterior variance is Λ^-1^*_i_*, with numerical stabilization. RNA/ADT decoders use count likelihoods and ATAC decoders use Bernoulli likelihoods.

The integration encoder used a shared backbone, 32 shared dimensions, 40 epochs and minibatches of 256. Reconstruction losses were normalized by feature count. Regularization weights were 0.001 for Kullback–Leibler divergence, 1 for bridge alignment, 0.1 for local geometry, 0.02 each for moving-average consistency and the gate prior, 0.05 for batch adversarial alignment, 0.1 for modality adversarial alignment and 0.5 for reference-distribution alignment. The variance-floor penalty had weight 0.2 and target 0.5; gate-monotonicity regularization was inactive. Bridge alignment compared paired modality coordinates, whereas the geometry term compared normalized pairwise distances in observed and latent spaces. Training seeds followed task configurations, without label-based checkpoint selection.

The spectral alternative reduced log-normalized RNA/ADT and binary TF–IDF ATAC blocks to 32 dimensions by truncated singular-value decomposition (SVD) (Halko et al. 2011). Standardized modality coordinates were orthogonally aligned where shared bridge cells were available, starting from RNA, and averaged across observed views. A fixed rule selected this alternative when final geometry loss exceeded three times its mean over epochs 20–30. Otherwise, the neural candidate underwent cluster-conditional batch refinement with 80 clusters, eight iterations, strength 1 and shrinkage 20. This rule selected 25 neural and five spectral routes; realized routes and feature counts are recorded per task.

### 4.3 Shared-feature semantic anchor

The semantic branch supplies a common gene-based reference across modalities. RNA genes were ranked by prevalence in RNA-observed cells, targeting 4,000 genes while retaining genes required by mapped proteins. ATAC peak centers were assigned to the nearest requested transcription start site within 100 kb, with weight exp(−*d/*50,000) for distance *d* in base pairs. Normalized protein names were matched to gene symbols. These mappings establish compatible coordinates rather than regulatory targets.

Each view was library-normalized to 10,000, log-transformed and scaled feature-wise without sparse-matrix centering. Stacked compatible views underwent truncated SVD to at most 32 dimensions, with padding when rank-limited, followed by separate standardization and centering of modality views. Each cell received its RNA view when observed, otherwise another available semantic view. Modalities with fewer than 64 compatible features could instead align a full-block spectral representation to a valid semantic view using at least two bridge cells (Schonemann 1966). For alignment weighting, log library size and log detected-feature count were scaled between their positive-value 10th and 90th percentiles; their geometric mean was clipped to [0.05, 1].

### 4.4 Adaptive semantic protection and alignment

For standardized semantic and candidate matrices *S* and *N* in matched cell order, orthogonal Procrustes alignment minimizes ∥*NQ* − *S*∥*_F_*, giving Ñ = *NQ*. Within each observation group, *k*nearest neighbors (*k* ≤ 20, excluding self) define 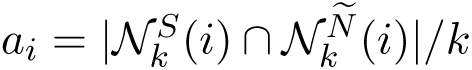 and median semantic-neighbor distance *d_i_*. With displacement 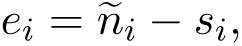, protection gives

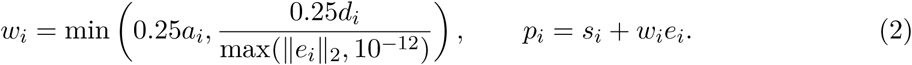

Thus, agreement determines the retained candidate displacement, subject to ∥*p_i_* − *s_i_*∥_2_ ≤ 0.25*d_i_*.

Adaptive mixing responds to observation-group separability in the candidate. Let *o* be the mean same-group fraction among 30 neighbors and 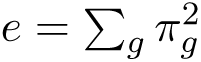 its chance expectation for group proportions *π_g_*. We compute

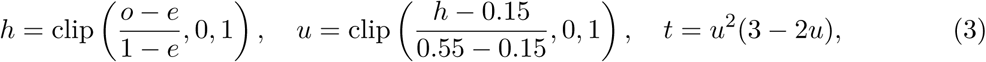

and return the mixture to the candidate coordinate frame:

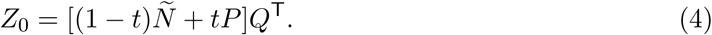

Zero intervention preserves the candidate route; greater group separation increases intervention. Neither calculation uses biological labels.

Alignment then applies three modality mutual-nearest-neighbor passes, one conditional location/scale step, batch mutual-nearest-neighbor alignment and two batch conditional location/scale iterations. Settings were 20 alignment neighbors, 40 smoothing neighbors, 40 conditional clusters and shrinkage 20; modality conditional strength was 1 + *t*. Local cross-group offsets were smoothed and cluster-specific group moments shrunk towards pooled estimates. The displacement bound applies to *P*, before these transformations. Existing component comparisons retain unprotected fusion, the semantic reference, a fixed 25% candidate residual, local protection without adaptive mixing and the full strategy. Their different coordinate and intervention handling makes them comparisons of complete strategies.

### 4.5 Molecular point recovery

Recovery used modality encoders with hidden widths 512 and 256, 32 shared and two additional latent dimensions, and gated product-of-experts fusion. Initial training used native Poisson RNA/ADT decoders for 100 epochs, batch size 256 and learning rate 2 × 10*^−^*^4^. Regularization weights were 0.001 for KL divergence, 1 for bridge alignment, 0.1 for geometry and 0.02 each for moving-average consistency and the gate prior; moving-average decay was 0.995.

Target heads transformed training-standardized source coordinates (standard-deviation floor 10*^−^*^4^) through two 256-unit Mish layers, with layer normalization after the first. Predictions were nonnegative for RNA/ADT and sigmoid-transformed for ATAC. Output biases were initialized to training means or their logits. Heads minimized mean squared error for RNA/ADT and binary cross-entropy for ATAC using AdamW, learning rate 10*^−^*^3^, weight decay 10*^−^*^4^, batch size 256 and gradient clipping at 2. Selection loss was assessed every ten epochs within a 100-epoch budget.

Joint adaptation updated encoders, gates and heads while cycling through both single-modality inputs and the paired input. A latent-anchor penalty of 0.01 constrained departure from the initial representation. Encoder/head learning rates were 5 × 10*^−^*^5^/2 × 10*^−^*^4^ for 50 epochs, then 10*^−^*^5^/5 × 10*^−^*^5^ for 20 refinement epochs. Refinement used ATAC loss weight 3 in RNA/ATAC tasks and selection checks each epoch. No auxiliary correlation loss or count thinning was used. Predictions from seeds 42, 0 and 1 were averaged equally on the target scale.

### 4.6 Conditional RNA residual diffusion

Diffusion conditions on the fixed three-member RNA point mean *y*(*c*) and the seed-42 member’s ATAC posterior statistics. The 34-dimensional posterior mean and log variance, the latter clipped to [−12, 12], were concatenated and standardized using training statistics with standard-deviation floor 10*^−^*^4^ to form *c*. Queries used ATAC alone. For feature *j*, residuals were 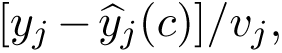 where *v_j_* is the training residual root mean square floored at 0.001. Conditioning concatenated *c* with 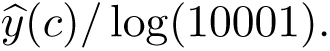 At timestep *τ*,

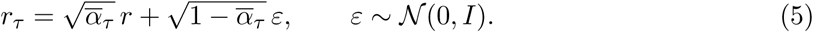

The denoiser predicted *ε* by mean squared error (Ho et al. 2020). It combined the residual, condition and a 32-dimensional sinusoidal time embedding in two 256-unit SiLU layers, with layer normalization after the first. A 100-step cosine schedule used offset 0.008 and noise variances clipped to [10*^−^*^4^, 0.999] (Nichol and Dhariwal 2021). Timesteps were sampled uniformly. AdamW used learning rate 2 × 10*^−^*^4^, weight decay 10*^−^*^4^, batch size 128 and gradient clipping at 2. Sampling traversed all 100 reverse steps, bounded clean-state predictions to the RNA target range and generated 32 samples per cell.

The conditional stochastic-head comparator shared the condition, anchor, hidden widths, optimizer and budget. Per feature, it predicted positivity probability, a softplus Gaussian location and log standard deviation clipped to [−4, 2]. Its loss combined positivity binary cross-entropy with a Gaussian surrogate on positive targets. Each family used seeds 42, 0 and 1, 100 epochs and the final checkpoint, without distribution-model configuration search.

Both families applied bounded mean projection: for each cell-feature pair, a common offset was added to all 32 samples before clipping to [0, log(10001)]. Thirty-two bisection iterations matched their mean to the fixed anchor within 2 × 10*^−^*^5^. This preserves the point prediction while retaining an empirical distribution. Historical Gaussian-residual and resampled-residual controls are reported separately.

### 4.7 Data sources and observation designs

All analyses reused public molecular measurements. Original studies, processed input locations and analysis subsets are listed in Supplemental Table S5. Integration comprised TEA PBMC RNA/ATAC/ADT (GSE158013) (Swanson et al. 2021), BMMC CITE-seq/Multiome (GSE194122), retinal Multiome (GSE196235) and bone marrow Ab-seq. Two TEA scenarios, two BMMC scenarios, Retina and Ab-seq contributed five configurations each (seeds 2024–2028). Recorded masks and batch roles defined observed inputs; common cells, feature vocabularies and annotations were retained without additional method-specific cell filtering. BMMC cell sampling and repeated cell instances followed each scenario definition.

Point recovery used PBMC-10x, Chen-2019 (mouse cortex, SNARE-seq; GSE126074) (Chen et al. 2019), Ma-2020 (mouse skin, SHARE-seq; GSE140203) (Ma et al. 2020) and the RNA/ADT subset of DOGMA (GSE166188) (Mimitou et al. 2021). PBMC-10x comprised 10,032 matched RNA/ATAC profiles from a healthy-donor PBMC sample profiled with 10x Genomics Multiome. We used the package-provided input imported in R with SingleCellMultiModal::scMultiome(“pbmc_10x”) (Eckenrode et al. 2023; 10x Genomics 2020). DOGMA retained 3,000 paired cells, four processing/condition batches and 210 shared ADT features. Donor-separated PBMC evaluation used GSE297529; pre-fit exclusion of GSM8994068 for invalid sparse indices left 16 donors and 35,422 selected cells. Human cortical matrices, annotations and fragments came from the BICCN challenge (BRAIN Initiative Cell Census Network (BICCN) 2021; Hodge et al. 2019); human–mouse motor-cortex inputs originated from NeMO SNARE-seq and GSE229169 Multiome.

### 4.8 Feature processing and recovery partitions

Integration selected up to 4,000 RNA, 10,000 ATAC and 256 ADT features by prevalence in observed cells. Neural RNA/ADT inputs were normalized to the median observed library size and log-transformed; ATAC was binarized. Recovery instead selected up to 2,000 RNA, 5,000 ATAC and 256 shared ADT features by training prevalence, excluding training-unobserved features. Its encoders received log-transformed raw RNA/ADT counts or binary ATAC. RNA/ADT targets were *y* = log[1 + 10^4^*x/* max(*L, ɛ*)], with *L* the selected-feature count total; ATAC targets were binary. Published feature vocabularies could reflect whole-cohort preprocessing, whereas recovery feature selection and coordinate standardization used training observations.

Development partitions used seed 913. With at least three donor/batch groups, one permuted group supplied testing, one calibration and the remainder training; calibration was divided into selection, policy and interval subsets. Otherwise, cells were split 60%/10%/10%/10%/10% into those five roles. PBMC-10x and Chen-2019 used cell splits. Ma-2020 trained on batches 56/53, calibrated on 55 and tested on 54; DOGMA used LLL_STIM/DIG_CTRL, LLL_CTRL and DIG_STIM, respectively. Fig. 4 development endpoints use selection partitions. Partition sizes and complete donor assignments appear in Supplemental Table S7.

GSE297529 separated donors across training, selection, policy fitting, risk calibration, interval calibration and testing. Up to 3,000 sorted-barcode cells per donor were sampled with seed 913 before fitting. Test donors GSM8994061/GSM8994074/GSM8994075 contributed 3,000/203/3,000 cells; selection used GSM8994069. RNA/ATAC were barcode-matched, fragments quantified on a fixed common region vocabulary and feature prevalence estimated in training donors.

Diffusion used unique paired cells from four TEA, three BMMC Multiome and eight Retina libraries, deduplicating scenario copies by original library/barcode and excluding BMMC CITE libraries. Within each library, seed 913 assigned 60%/20%/20% to training/validation/evaluation, yielding 5,105/3,449/10,065 evaluation cells by source (Supplemental Table S3). Training prevalence selected 2,000 RNA and 5,000 ATAC features. Three point members per source used the recovery schedule with validation-based checkpoint selection; evaluation RNA entered neither fitting nor selection. These within-library holdouts are distinct from development point-recovery and donor-separated partitions.

### 4.9 Comparator implementations

Integration compared Palette, MIDAS, scMoMaT, StabMap and MoNAE (Sheng et al. 2026; He et al. 2024; Zhang et al. 2023; Ghazanfar et al. 2024; Tang et al. 2024), using method-specific preprocessing and budgets (Supplemental Table S6). Evaluation labels did not guide checkpoint selection.

For recovery, Palette transferred measured training features, whereas StabMap, scMoMaT and MoNAE representations used the common molecular heads described above; a separate MoTRUST representation-plus-head control used the same readout. These adapters fitted paired training cells together with source-only queries, a transductive setting distinct from fixed-model inference for MoTRUST and MIDAS. Query targets and labels were excluded. Two directions and three seeds across four development datasets and GSE297529 yielded 120 adapter fits; predictions were averaged before scoring. Detailed settings are given in Supplemental Methods.

An empty affinity graph in all three Chen-2019 ATAC-to-RNA Palette fits required joint scaled source principal components before feature transfer, without query targets. Development MIDAS endpoints use the strongest previously evaluated native, matched-head or calibrated variant per endpoint; GSE297529 reports native and matched-head three-member ensembles separately. Endpoint-specific MIDAS variants are listed in Supplemental Table S9.

### 4.10 Evaluation metrics and statistical summaries

Biological conservation averaged NMI, ARI, cASW and cLISI; batch correction averaged batch silhouette, iLISI, neighborhood batch acceptance and graph connectivity (Hubert and Arabie 1985; Rousseeuw 1987; Luecken et al. 2022; Buttner et al. 2019). All components were required to be finite, and

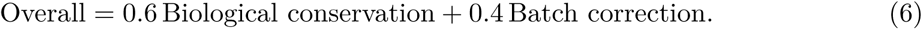

Cell-type-balanced modality mixing was evaluated separately. The fixed evaluator and component transformations are specified in Supplemental Methods.

Point recovery used RMSE, feature-wise Pearson correlation and ATAC AUPRC (Saito and Rehmsmeier 2015). Fig. 4 averages RMSE/Pearson equally over directions; AUPRC uses ATAC targets, excluding DOGMA. GSE297529 summaries weight the three test donors equally.

For *K* predictive samples and observed target *y*, marginal CRPS was

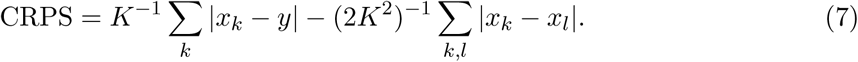

CRPS was averaged over evaluation cells/features, then seeds; relative changes were ratios of method-level seed means minus one. CRPS and energy score retained the *K*^2^ denominator because projection couples samples (Gneiting and Raftery 2007). Energy score, interval coverage/width and post-fit strata are detailed in Supplemental Methods.

Paired one-sided Wilcoxon tests used Holm adjustment (Wilcoxon 1945). Tests and standard errors were descriptive: task configurations and donor-role permutations reused cells, while training seeds and cross-seed pairings shared evaluation partitions. Dataset-block summaries accompanied task comparisons. Weight sensitivity reused embeddings with biological weights 0.40–0.80 in 0.05 increments.

### 4.11 Donor uncertainty calibration

The earlier donor analysis calibrated point-ensemble outputs separately from diffusion intervals (Supplemental Fig. S2). A feature-scale network combined seven source/prediction summaries and 32 training-fitted source SVD coordinates in a 64-unit SiLU layer, 16 tanh factors, feature loadings and a feature-specific slope on standardized predictions. Log-variance adjustments were clipped to [−4, 4] around policy-fit residual variance. Training used policy-fit donors only, Gaussian residual loss, AdamW learning rate/weight decay 10*^−^*^3^, batch size 256, 100 epochs and seed 42. Interval-calibration quantiles of |*y* − *ŷ*|*/σ* defined clipped *ŷ* ± *qσ* intervals at coverage 0.90, using quantile rank ⌈(*n* + 1)0.90⌉*/n*. ATAC class sets separately calibrated *p̂* among zeros and 1 − *p̂* among ones, retaining classes with fewer than ten calibration examples. These partitions do not establish distribution-free coverage under donor shift (Lei et al. 2018).

### 4.12 Cortical and cross-species analyses

BICCN contained 31,960 paired identifiers. Requiring positive counts in both assays and sampling 3,000 cells per donor with seed 913 yielded 9,000 cells. NeuN-enriched and repeat libraries were grouped by their three recorded donors. All six permutations assigned one bridge, one RNA-only query and one ATAC-only query donor. Subclass labels defined evaluation cell types; class/cluster labels supported secondary summaries. Fifteen-bridge-neighbor majority vote supplied query macro-F1; rare bridge-supported types were the lower quartile of positive bridge counts.

Fragments were reconstructed on 616,439 fixed peaks using barcode-matched insertions at start and end minus one under half-open BED coordinates, ignoring PCR multiplicity and capping cell-peak counts at four. RNA/ATAC signals were log(1 + 10^4^*x/L*) with respective gene/peak totals. Gene-local accessibility used 2-kb and 100-kb transcription-start-site windows for PVALB, SST, VIP, LAMP5, GAD1, GAD2, CUX2 and TLE4; marker AUROC measured designated-subtype separation. Hidden RNA was the mean measured RNA of 15 bridge neighbors, independently of recovery heads and diffusion. Inhibitory-marker Spearman correlations were summarized with equal donor weights. The axis [*z*(LHX6) + *z*(SOX6) − *z*(PROX1) − *z*(NR2F2)]*/*2 used bridge-inhibitory RNA for predictive standardization. Residual associations adjusted for subtype/depth, with 999 within-subtype permutations and Benjamini–Hochberg correction over 48 comparisons (Benjamini and Hochberg 1995); they do not establish causal regulation.

Cross-species evaluation sampled 3,000 paired cells per species with seed 2024, retaining 4,659 identically ordered homologous RNA groups and 5,000 prevalence-selected peaks per species. Shared normalized RNA supplied the semantic anchor because homolog-group identifiers lacked gene/TSS mapping; separate Bernoulli heads decoded species-specific ATAC. Comparator adapters retained this arrangement and recorded budgets. Palette used normalized-data singular-value initialization for single-assay ATAC blocks skipped by its original routine, retaining shared-RNA Bi-sPCA and subsequent inference. Bidirectional five-neighbor voting in standardized embeddings used fixed true target-label sets for macro-F1; absent and rare supported types were summarized separately. The RNA-anchor control used identical cells. Species and assay were confounded, and ATAC-only transfer was not evaluated.

### Software availability

The MoTRUST source code is available at https://github.com/BryantLuffy/MoTRUST.

## Data access

This study reanalyzed previously published datasets and generated no new primary sequencing data. Original study accessions and public input sources are listed in Supplemental Table S5. They include GEO accessions GSE158013, GSE194122, GSE196235, GSE126074, GSE140203, GSE166188, GSE297529 and GSE229169, together with the cited Figshare, 10x Genomics, BICCN and NeMO resources. For GSE297529, the analyses used public processed matrices and supplied fragments; raw sequencing reads are not openly provided by the original study.

## Author contributions

Qingchun Liu and Yan Xu conceived the study. Qingchun Liu developed the methodology and software, performed the data analyses, and wrote the original draft. Yan Xu supervised the research, acquired funding, and reviewed and edited the manuscript. Both authors read and approved the final manuscript.

## Ethics and consent

This study involved secondary analysis of previously published datasets and did not involve new participant recruitment or new sample collection. The original studies are cited in Methods and Supplemental Table S5.

